# When does more data help? Spectral Geometry and Scaling Laws in MRI Transformers

**DOI:** 10.64898/2026.07.14.738571

**Authors:** Tamoghna Chattopadhyay, Kartik Shelar, Sophia Thomopoulos, Paul M. Thompson

**Affiliations:** Imaging Genetics Center, Mark and Mary Stevens Neuroimaging and Informatics Institute, Keck School of Medicine,University of Southern California, Marina del Rey, CA, United States

**Keywords:** Vision Transformer, Spectral Concentration, Scaling Laws, Zeta Law, Self-Supervised Learning, Mahalanobis Distance

## Abstract

Scaling laws describe how model performance improves as the amount of training data increases, and recent theories such as the zeta law suggest that scaling behavior is influenced by the eigenspectrum of the model’s latent representation. Here, we evaluated whether the distribution of discriminative signals across spectral modes predicts the future scaling behavior, for MRI transformers trained for disease classification. We trained three supervised 3D vision transformers (ViT3D, MINiT, and NIT) for Alzheimer’s disease classification using 2,822 training scans from the Alzheimer’s Disease Neuroimaging Initiative (ADNI); we compared their encoder spectra with that of a frozen self-supervised DINO ViT-B/16 encoder adapted to 3D MRI. The supervised models learned highly concentrated representations, with 90–96% of CLS-token variance captured by a single principal component, whereas DINO distributed signal across many latent directions. Via spectral expansion of the Mahalanobis signal, we found that supervised training concentrated disease information into a single dominant mode, while self-supervised training produced a richer spectral geometry with higher effective rank and discoverability. This led to different scaling behavior: supervised models exhibited flatter AUC(*N*) curves, yet DINO continued to improve as sample size increased, gaining 11.0 percentage points from N=50 to N=2,822. Overall, the spectral distribution of the discriminative signal, for these different encoder types, influenced how much performance remained discoverable as sample size increased. Distributed representations may retain signal across many latent modes and continue to improve with additional data, whereas concentrated representations tend to exhaust most of the discoverable signal at much lower sample sizes.

## 1 Introduction

Vision transformers (ViTs) are now a dominant architecture for image recognition, with increasing success in medical imaging, where 3D variants are now applied to volumetric brain MRI for clinical tasks such as disease classification [1,2]. In neuroimaging, a key application is the automated discrimination of cognitively normal (CN) individuals from those with neurodegenerative diseases such as Alzheimer’s disease and Parkinson’s disease. using multimodal images, such as structural and diffusion MRI, and amyloid or tau PET. This task, along with related tasks such as predicting future clinical decline [3,4] and neuropathology classification from *in vivo* MRI [5] have direct clinical relevance for early diagnosis and patient stratification for drug trials and genetic analysis of underlying disease mechanisms [6-8]. Despite strong performance, transformer-based classifiers often show diminishing returns as training set size grows beyond the low-thousands, and neuroimaging studies frequently attribute this limitation to data scarcity [9,10]. The spectral geometry of learned representations provides a principled framework to examine this scaling failure. Recent theoretical work on the zeta law [11,12] proposes that the data-scaling behaviour of a classifier is influenced by the eigenspectrum of its encoder’s embedding space. As the training sample size, *N*, increases, the application of Vershynin’s [13] concentration inequality, Weyl’s inequality and the Davis-Kahan principle imply that more modes, *K(N)*, of the data covariance can be reliably estimated [12]. This eigenbasis can then be used for spectral decomposition of the class difference vector in the embedding space, and the Mahalanobis energy of the discriminative signal builds up over an increasing set of eigenvectors as the training *N* increases [11,12]. The framework predicts that encoders with steep eigenvalue spectra, where one (or a few) principal components capture the vast majority of explained variance, will be less sensitive to dataset size beyond a certain point, regardless of how much data is collected. Spectral collapse, whereby some neural networks concentrate variance into very few dimensions, has been documented in self-supervised learning [14,15] and linked to the training objective design. Contrastive and self-supervised objectives that enforce consistency across multiple augmented views discourage such spectral collapse by requiring the encoder to represent many orthogonal sources of variation in embedding space [16,17]. Supervised objectives, by contrast, optimize directly for a single binary decision boundary and have no such distributional pressure. The impact of spectral collapse on data-scaling behavior has not been systematically studied in medical imaging, where supervised models are often trained on only a few thousand labeled samples.

Here we investigate whether spectral collapse limits data scaling in 3D MRI transformers, and whether the training objective rather than architecture is its primary cause. We train three supervised transformers (ViT3D, MINiT, and NIT [1,2]) on ADNI brain MRI data [18] for CN-versus-dementia classification, a representative benchmark expected to generalize to other tasks. We then analyze their CLS-token embeddings using a Mahalanobis signal framework [11,12]. We also evaluate a 3D-inflated DINO ViT-B/16 encoder [19], freezing its self-supervised ImageNet-1k backbone and training only a linear classifier. Comparing spectral metrics across all four models tests whether spectral collapse is driven primarily by the training objective rather than architecture, with DINO providing a positive control.

## 2 Theoretical Background

### Eigenvalue Spectrum and Spectral Slope

Let *E* ∈ *R* ^{*NxD*}^ be mean-centred CLS embeddings [20], where *D* is the dimension of the embedding space (encoder) and N is the number of samples mapped through the trained encoder. In the theoretical limit where N is sufficiently high, the covariance eigensystem yields eigenvalues λ_1_ ≥ λ_2_ ≥ … ≥ λ_*D*_, although only the top *K(N)* of these can be reliably estimated from a finite data sample of size *N* [12] We model the leading spectrum as a power law, 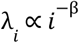, where we estimate the spectral slope β > 0 by OLS regression of log(λ _*i*_ ) on log(*i*) over the leading 50 modes, which consistently fell within the stable log-linear spectral regime across all models. Complementary summary statistics that describe the encoder spectrum are the *effective rank*, 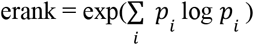 with 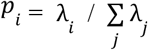, and 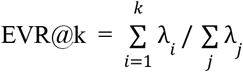 . High spectral slope, β, implies that the variance of the data is concentrated in few leading components; low β implies a richer, more isotropic embedding space where more components are needed to fully describe the data.

### Mahalanobis Signal and Admission Thresholds

Let **d** = **μ**_**{dem}**_ **− μ**_**{cn}**_ be the class-mean difference in embedding space, with projections 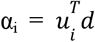, onto each eigenmode *i*. Direction *i* contributes a net positive signal when 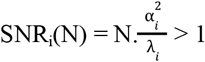, which first holds at the *admission threshold* 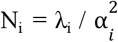 . The squared Mahalanobis distance, summed over SNR-admissible directions, is 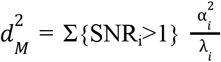, giving predicted AUROC AUC_pred_ = Φ(d_M_ / √2) under Gaussian class-conditionals [12]. The rate of improvement depends on how discriminative signal is distributed across the spectrum: concentrated representations tend to saturate early, whereas distributed representations keep gaining discriminative power as additional modes become discoverable. Thus, AUC(*N*) scaling reflects how quickly additional spectral modes are revealed as *N* grows. Under power-law spectra, 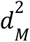 is governed by a partial sum of the Riemann zeta function, ζ_K(*N*)_ (q) = Σ{i=1 to K(*N*)}*i* _−β_, so that AUC(*N*)≈Φ(√ [(C/2) . ζ_K(*N*)_(q)] ), Because classification performance is determined by the growth of this truncated zeta sum as the discoverability horizon expands with sample size, we term this relationship the ***zeta law of generalization*** [12].

### L(k) Scaling Exponent and Error Floor

Define *L*(*k*) = 1 − AUROC(*k*) where AUROC(*k*) is measured by a logistic probe on the top-*k* PCA projection of the training embeddings. We fit *L*(*k*) = *A*/*k*^α^ + δ. The exponent α encodes how rapidly additional PC dimensions improve classification: α = 0 indicates spectral collapse, where all discriminant signal is concentrated in PC1. The constant δ is the irreducible error floor, set by the encoder’s total discriminative capacity and its match to the task domain [11].

## 3 Methods

### Dataset

We analyzed publicly available ADNI data [18], selecting cognitively normal (CN) and dementia subjects. This well-studied benchmark was chosen to illustrate the concepts and is known to yield strong classification performance. Subjects were split 70/15/15 at the subject level, stratified by diagnosis and sex, yielding 2,822 training scans (811 subjects), 576 validation scans (174 subjects), and 587 test scans (174 subjects). All inputs were 3D T1-weighted MRI volumes (96×112×96 voxels). To isolate encoder effects, no tabular covariates were used, although age and demographic variables are commonly included in practice [21].

### Supervised Architectures

Three supervised 3D transformer architectures (**Table 1**) were trained end-to-end with a Linear (256→1) and sigmoid classification head on the CLS token. Training parameters were AdamW (learning rate: 1×10^−4^, cosine annealing, 5-epoch warmup), weight decay 0.01, FP16 AMP, batch 16, 100 epochs, early stopping (patience 15, val AUC), class-weighted BCE (pos_weight=1.4391).

**Table 1.** Supervised Architecture Summary.

| Model | Architecture | Layers / Heads | Dim. | Param. |
| --- | --- | --- | --- | --- |
| VIT-b/16 | Global ViT, $16^3$ patches, 252 tokens + CLS | 6 / 8 | 256 | 5.85M |
| MINIT | Block encoder (4L) + ABMIL aggregation | 4 / 8 | 256 | 3.27M |
| NIT | Global ViT with NIT attention blocks | 6 / 8 | 256 | 5.85M |

### DINO ViT-B/16 Positive Control

To test whether spectral collapse is caused by the training objective or the transformer architecture, we adapted DINO ViT-B/16 [19], pre-trained on ImageNet-1k (∼1.28M images) by enforcing CLS token consistency across augmented views to 3D MRI. The 3D inflation comprises four steps: (1) the Conv2d patch embedding (768, 3, 16, 16) was adapted to Conv3d(1, 768, 16) by averaging RGB input channels and replicating the 2D kernel T = 16 times along the depth axis, divided by T to preserve activation magnitude; (2) the CLS token is transferred directly; (3) positional embeddings are set to zero (the 2D 14×14 grid has no canonical interpolation to a 3D 6×7×6 grid); (4) all 12 transformer blocks are transferred and frozen. The backbone processes 252 patches in a (B, 253, 768) sequence, structurally identical to ViT3D. A fresh Linear(768→1) head (769 parameters) is the only trainable component, trained for 30 epochs (AdamW, lr = 1×10^−3^, cosine). Zeroing positional embeddings may reduce spatial discriminability, but is unlikely to explain the distributed spectral geometry observed, as it affects all directions equally and would not prevent spectral collapse.

### Comparability caveat

DINO ViT-B differs in embedding dimension (D=768 vs. 256) and parameter count (88.4M vs. 3.27–5.85M). All spectral comparisons use scale-free metrics (β, erank, EVR@k, α, δ) insensitive to absolute embedding scale. The monotone ordering across all four metrics across all four models supports training objective as the dominant driver, though architecture and capacity cannot be fully excluded without a matched supervised ablation.

### Spectral Analysis

CLS embeddings were extracted in evaluation mode from each model’s best checkpoint and all spectral analyses were fitted on training-split embeddings only. Four pipelines were applied identically to all four models: OLS regression of log(λ_i_) on log(*i*) over *i* = 1…50 yields β, erank, EVR@k; Mahalanobis spectrum α = u ^T d, SNR-truncated d_M_, admission thresholds 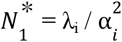, and PC1 signal concentration; L(*k*) = 1 − AUROC(k) power-law fit over k = 1–80 yields (α, δ); AUC(*N*) subsampling sweep over N ∈ {50, 100, 200, 400, 800, 1600, 2822}, 5–10 seeds, stratified by diagnosis.

## 4 Results

### Classification Performance

Table 2 reports test-set performance. Supervised models achieved test AUC 0.818–0.878. DINO achieved 0.791 with only 769 trainable parameters. The larger val-to-test gap for DINO (0.863 → 0.791) versus supervised models (≤0.029) reflects ImageNet-to-MRI domain shift in the frozen backbone. This is expected and is explained in Sections 4.4 and 5.2.

**Table 2.**
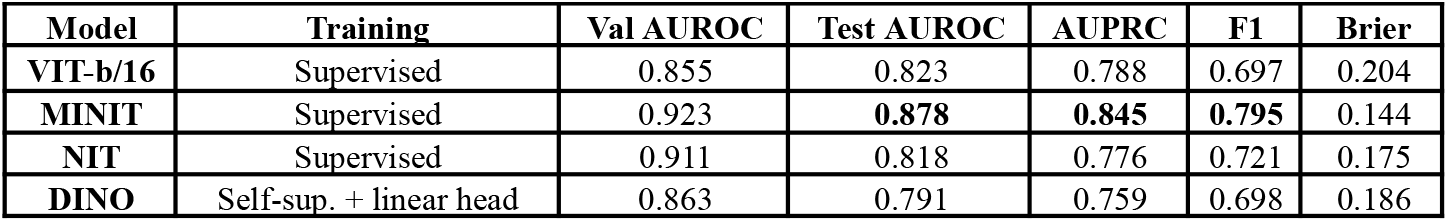
Supervised Architecture Summary.

### Eigenvalue Spectra and Spectral Slope β

Table 3 reports spectral geometry metrics. The spectral slopes form a monotone ordering consistent with the supervision gradient, though embedding dimension and model capacity also differ across models and may contribute to this pattern: ViT3D (2.311) > MINiT (2.271) > NIT (2.002) > DINO (1.148). All supervised models have EVR@1 = 0.897–0.955: a single PC captures 90–96% of total embedding variance. DINO has EVR@1 ≈ 0.19 - no collapse. Effective rank rises from 1.27–1.72 (supervised) to 52.18 (DINO), a 30-40× increase. **Fig. 1** overlays the four eigenvalue log-log plots.

**Table 3.** Spectral slope β, effective rank, participation ratio PR, and EVR at three cutoffs. DINO R^2^ not reported; power-law fit approximate for richer spectra; β = 1.148 is a robust fit.

| Model | $\beta$ | $R^2$ | erank | PR | $\text{EVR}@1$ | $\text{EVR}@10$ | $\text{EVR}@50$ |
| --- | --- | --- | --- | --- | --- | --- | --- |
| VIT-b/16 (Supervised) | 2.311 | 0.952 | 1.27 | 1.10 | 0.955 | 0.998 | 1.000 |
| MINIT (Supervised) | 2.271 | 0.955 | 1.38 | 1.13 | 0.941 | 0.996 | 0.999 |
| NIT (Supervised) | 2.002 | 0.959 | 1.72 | 1.24 | 0.897 | 0.988 | 0.997 |
| DINO (Self-Supervised) | 1.148 | — | 52.18 | 18.66 | $\approx 0.19$ | $\approx 0.50$ | $\approx 0.85$ |

**Fig. 1.**
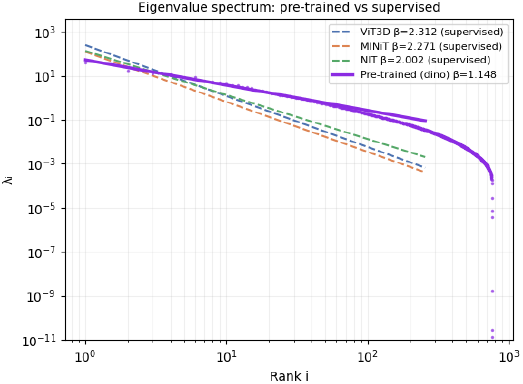
Log-log eigenvalue spectra for all four models. Spectral slope β (OLS over i = 1…50) is shown in the legend. Supervised models cluster between β = 2.002 and 2.312. DINO ViT-B/16 (β = 1.148) shows slower variance decay, distributing variance across more principal components.

### Mahalanobis Spectrum and Signal Concentration

Table 4 reports the Mahalanobis spectrum. Three metrics follow the monotone supervision gradient.

**Table 4.** Mahalanobis spectrum. 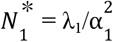 is the PC1 admission threshold; 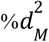 is the fraction of total Mahalanobis signal in PC1.

| Model | $k_{\text{snr}}$ | $d_M$ | $\vert \alpha_1\vert $ | $\vert \alpha_2\vert $ | $\vert \alpha_1\vert /\vert \alpha_2\vert $ | $N_1^*$ | $\%d_M^2$ PC1 |
| --- | --- | --- | --- | --- | --- | --- | --- |
| VIT-b/16 (Supervised) | 66 | 1.872 | 16.932 | 0.512 | 33× | 0.3 | 90.6% |
| MINiT (Supervised) | 71 | 1.753 | 6.711 | 0.022 | 305× | 0.4 | 91.2% |
| NIT (Supervised) | 71 | 1.591 | 6.264 | 0.243 | 26× | 0.5 | 79.8% |
| DINO (Self-Supervised) | 79 | 1.318 | 0.210 | 0.055 | 3.8× | 2.93 | 31.0% |

#### PC1 dominance ratio

|α_1_|/|α_2_| drops from 26–305× for supervised models to 3.8× for DINO. For MINiT, |α_1_| = 6.711 vs. |α_2_| = 0.022 (305×): essentially all Mahalanobis signal resides in a single direction. DINO’s signal is genuinely distributed across PC directions.

#### Admission threshold

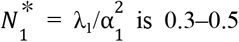 for supervised models so the leading discriminative mode is available from the outset. In contrast, DINO has 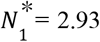 requiring roughly three training samples before PC1 becomes admissible.

#### Signal concentration

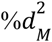 in PC1 drops from 79.8–91.2% (supervised) to 31.0% (DINO). Total d_M_ is lower for DINO (1.318 vs. 1.591–1.872) because DINO was not trained to maximize CN vs. dementia separation. The contrast is distributional, not in total magnitude. **Fig. 2** shows the Mahalanobis spectrum 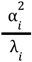 for all four models.

**Fig. 2.**
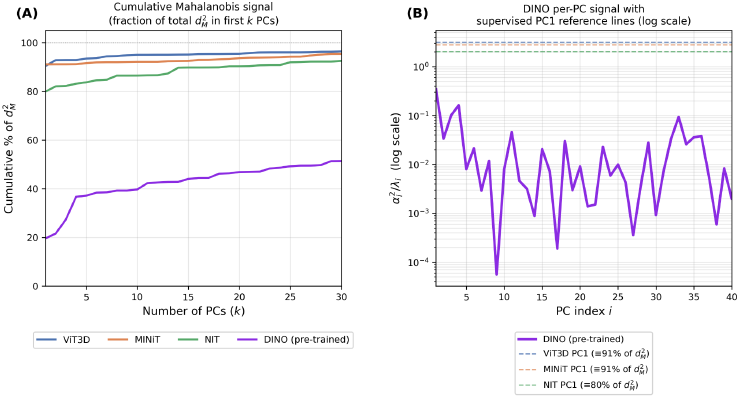
Mahalanobis signal distribution across principal components for all four models. (A) Cumulative fraction of total 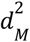 captured by the first *k* PCs. Supervised models (ViT3D, MINiT, NIT) accumulate 80–91% of their total Mahalanobis signal in PC1 alone and quickly plateau, indicating that the discriminative signal is concentrated in a single direction. DINO starts at ∼20% at k=1 and rises gradually, reaching only ∼50% by k=30, confirming that its signal is distributed across many directions. (B) Per-PC Mahalanobis signal 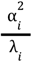 for DINO (log scale), with the PC1 signal levels of the three supervised models (*dashed horizontal lines*). The supervised PC1 values sit 1–2 orders of magnitude above DINO’s individual PC contributions, yet DINO’s signal accumulates across many PCs while the supervised models contribute almost nothing beyond PC1.

### Four-Model Comparison

Table 5 summarizes all spectral metrics. The one exception to the monotone ordering is δ, addressed below.

**Table 5.** Four-model spectral comparison. Δ = AUC_obs − AUC_pred (geometry-AUC gap). Monotone direction: β ↓, erank ↑, α ↑ along supervision gradient.

| Model | $\beta$ | erank | $d_M$ | $\alpha$ | $\delta$ | $\Delta$ | AUC obs |
| --- | --- | --- | --- | --- | --- | --- | --- |
| VIT-b/16 (Supervised) | 2.311 | 1.27 | 1.872 | 0.000 | 0.189 | -0.085 | 0.823 |
| MINIT (Supervised) | 2.271 | 1.38 | 1.753 | 0.000 | 0.126 | -0.014 | 0.878 |
| NIT (Supervised) | 2.002 | 1.72 | 1.591 | 0.119 | 0.090 | -0.051 | 0.818 |
| DINO (Self-Supervised) | 1.148 | 52.18 | 1.318 | 0.802 | 0.214 | -0.034 | 0.791 |

#### L(k) scaling exponent α

α rises 0.000 → 0.000 → 0.119 → 0.802. For ViT3D and MINiT, *L*(*k*) is completely flat: adding PC dimensions provides zero benefit;discriminant signal is heavily concentrated in PC1. For DINO, L(k) falls from 0.345 at k = 1 to 0.207 at k = 80, a 40% error reduction as more PC dimensions are included.

#### Error floor δ

δ_{DINO} = 0.214 exceeds δ_{NIT} = 0.090, reflecting ImageNet-to-MRI domain mismatch rather than geometric failure: DINO’s features encode generic visual structure rather than disease-specific anatomy, and the lower d_M_ = 1.318 may reflect this task misalignment. The smaller geometry-AUC gap |Δ| = 0.034 (vs. 0.085 and 0.051 for ViT3D/NIT) confirms that distributed geometry generalises better despite lower total signal. Domain-matched self-supervised pre-training would eliminate this δ deficit.

### AUC(N) Scaling

Table 6 and **Fig 3** show the AUC(*N*) subsampling results. Supervised models are near-flat: ViT3D and MINiT show −0.7% and −0.4% change from N = 50 to N = 2822; NIT shows +3.4%. DINO shows +11.0 percentage points (0.683 → 0.793). This is a consequence of spectral collapse: when N_1_ < 1, performance saturates before N = 50; adding data provides no benefit. When N_1_ = 2.93, PC1 is not admitted until N ≈ 3 and performance rises across the full training range.

**Table 6.** AUC(N) subsampling sweep (5 seeds each for DINO; 10 seeds for supervised models).

| Model | AUC(N=50) | AUC(N=800) | AUC(N=2822) | Total gain |
| --- | --- | --- | --- | --- |
| VIT-b/16 (Supervised) | $\approx 0.815$ | $\approx 0.812$ | $\approx 0.808$ | -0.7 pp |
| MINIT (Supervised) | $\approx 0.878$ | $\approx 0.875$ | $\approx 0.874$ | -0.4 pp |
| NIT (Supervised) | $\approx 0.817$ | $\approx 0.837$ | $\approx 0.851$ | +3.4 pp |
| DINO (Self-Supervised) | 0.683 | 0.784 | 0.793 | +11.0 pp |

**Fig. 3.**
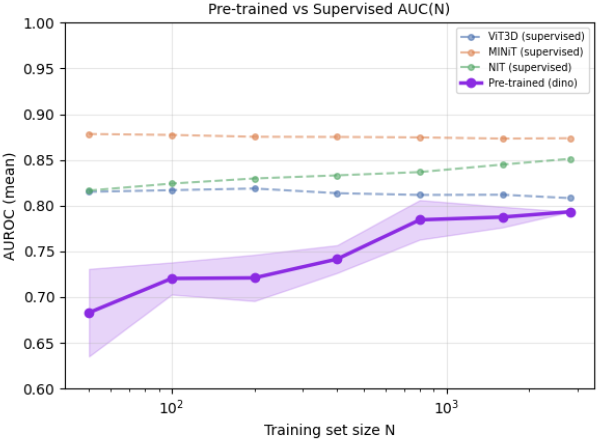
AUC(*N*) subsampling curves. Supervised models (ViT3D, MINiT, NIT) are flat across N = 50–2,822, consistent with N_1_ < 1. DINO (mean ± sd across 5 seeds, shaded) shows a rising curve from AUC = 0.683 at *N* = 50 to 0.793 at N = 2,822, consistent with the distributed Mahalanobis signal.

## 5 Discussion

The DINO results suggest that spectral collapse is driven primarily by the supervised training objective rather than architecture. While differences in model capacity and pre-training data cannot be fully excluded without a matched supervised ViT-B/16, the consistent ordering across four spectral metrics and AUC(*N*) provide convergent evidence for the training objective as the dominant factor. A promising remedy is domain-matched self-supervised pre-training on 3D brain MRI, which could combine distributed geometry (α > 0) with stronger signal (lower δ). DINO’s lower d_M = 1.318 likely reflects its encoding of generic visual structure rather than disease-specific anatomy. Rather than undermining the framework, this result points to missing domain alignment. DINO’s smaller |Δ| suggests that its distributed geometry generalizes better to unseen data, with the domain gap limiting signal rather than generalization. In the AUC(*N*) staircase framework [9], supervised models have N_1_ = 0.3–0.5, effectively collapsing the staircase, whereas DINO partially restores the observable regime (N_1_ = 2.93). Full recovery may require a domain-matched encoder with N_i > 50 across multiple directions, a directly testable hypothesis.

### Limitations

(1) DINO differs in dimension, capacity, and training corpus from supervised models - a supervised ViT-B/16 at D=768 would provide a stronger ablation; (2) 3D inflation zeros positional embeddings, limiting spatial encoding; (3) our analyses used a single dataset and binary task, so generalization to subtler problems (e.g., MCI conversion) or multisite problems is unknown; (4) the Gaussian equal-covariance AUC formula may not hold exactly for supervised embeddings; (5) confidence intervals are not reported for supervised AUC(*N*) curves. Future work will study the multisite case, where harmonization methods help to pool multiple datasets, and “double-zeta” or hypergeometric fractal laws could account for the spectral mixing of site, imaging protocol, and biological effects.

## 6 Conclusion

Spectral collapse - the concentration of CLS embedding variance into a single principal component - is the primary constraint on data scaling in supervised 3D MRI transformers and appears to be driven mainly by the training objective rather than architecture. This conclusion is supported by four independent spectral metrics and the observed AUC(*N*) behavior.. A DINO ViT-B/16 inflated to 3D shows β = 1.148, erank = 52.18, α = 0.802, and an actively rising AUC(*N*) curve (+11.0 pp), versus β = 2.00–2.31, erank ≤ 1.72, α = 0.000, and flat AUC(*N*) for all supervised counterparts. DINO’s higher error floor (δ = 0.214) likely reflects the ImageNet-to-MRI domain gap, motivating domain-matched self-supervised pre-training. Future work could test whether β, effective rank, and the discoverability horizon *K*(*N*) predict the value of additional training data before large-scale retraining.

## Acknowledgment

Supported by NIH grants U01 AG068057 and S10OD032285.

